# Fine-grained deconvolution of cell-type effects from human bulk brain data using a large single-nucleus RNA sequencing based reference panel

**DOI:** 10.1101/2022.06.23.497397

**Authors:** Edwin J.C.G. van den Oord, Karolina A. Aberg

## Abstract

Brain disorders are leading causes of disability worldwide. Gene expression studies provide promising opportunities to better understand their etiology. When studying bulk tissue, cellular diversity may cause many genes that are differentially expressed in cases and controls to remain undetected. Furthermore, identifying the specific cell-types from which association signals originate is key to formulating refined hypotheses of disease etiology, designing proper follow-up experiments and, eventually, developing novel clinical interventions. Cell-type effects can be deconvoluted statistically from bulk expression data using cell-type proportions estimated with the help of a reference panel. To create a fine-grained reference panel for the human prefrontal cortex, we analyzed data from the seven largest single nucleus RNA-seq (s_n_RNA-seq) studies. Seventeen cell-types were robustly detected across all seven studies. To estimate the cell-type proportions, we proposed an empirical Bayes estimator that is suitable for the new panel that involves multiple low abundant cell-types. Furthermore, to avoid the use of a very large reference panel and prevent challenges with public access of nuclei level data, our estimator uses a panel comprising mean expression levels rather than the nuclei level s_n_RNA-seq data. Evaluations show that our empirical Bayes estimator produces highly accurate and unbiased cell-type proportion estimates. Transcriptome-wide association studies performed with permuted bulk RNA-seq data showed that it is possible to perform TWASs for even the rarest cell-types without an increased risk of false positives. Furthermore, we determined that for optimal statistical power the best approach is to analyze all cell-types in the panel as opposed to grouping or omitting (rare) cell-types.

## Introduction

Brain disorders such as mood disorders, dementias, stress related disorders, neurodevelopmental disorders, seizure disorders, and addictions are leading causes of disability worldwide^1^. Gene expression studies provide promising opportunities to better understand their etiology. The human brain comprises multiple types of excitatory and inhibitory neurons as well as glia cells such as astrocytes, oligodendrocytes, and microglia^2-4^. As cells differ in their functions, gene expression will typically also vary across these cell-types. When studying bulk tissue, this cellular diversity may cause many genes that are differentially expressed in cases and controls to remain undetected^5^. That is, association signals will be “diluted” if they affect only one cell-type, may cancel out if they are of opposite signs across cell-types, and may be undetectable if they involve low-abundant cells. For example, even a very large expression difference of one standard deviation (SD) in a cell-type with a 1% frequency would be impossible to detect in bulk tissue as the expression difference would reduce to one-hundredth of a SD (i.e., 1%×1 SD = 0.01 SD).

Identifying the specific cell-types from which association signals originate is also critical for scientific progress and important from a translational perspective. First, it allows formulating refined hypotheses about disease etiology. For example, the involvement of microglia may point to disrupted immune response and neuroinflammation of the brain^6^, a loss of neuronal function may point to neurodegeneration^7^, and the involvement of the myelin-producing oligodendrocytes may suggest disrupted neuronal communication^8^. Second, knowledge about the cell-type is important to design proper *in vitro* or *in vivo* functional follow-up studies. Thus, as gene expression may only be altered in specific cells, such studies require the right choice of cultured cells or experimental tools (e.g., the use herpes simplex virus type 1 as a vector for locus-specific editing is of primary relevance for association findings in neurons^9,10^). Third, cell-type knowledge is key for developing novel and effective treatments. For example, drugs often work by interacting with receptors on the surface of cells. Receptor molecules have a specific three-dimensional structure, which allows only substances that fit precisely to attach to it. From a drug development perspective, designing drugs that interact specifically with receptors from particular cell-types is also highly desired since non-specific drugs can cause more side effects.

Single cell/nucleus RNA sequencing is a relatively new approach to study cellular diversity. In comparison to whole cells, nuclei are more resistant to mechanical assaults and are less vulnerable to the tissue dissociation process. This makes single nucleus RNA sequencing (s_n_RNA-seq) the more suitable option for frozen post-mortem brain tissue^11^. With this approach intact nuclei are first isolated and partitioned so that the content of each nucleus can be labeled with a unique identifier. A labeled sequencing library is subsequently generated and sequenced for each individual nucleus. Cell-type specific effects can also be deconvoluted statistically from bulk RNA-seq data^5,12^. Deconvolution was introduced 20 years ago^12^ and has been experimentally validated using, for instance, predesigned mixtures^13^. Deconvolution is most effective when performed with a reference panel^14^, typically generated from expression profiles of the cell-types present in the target tissue from a small number of reference samples. The reference panel is used to estimate cell-type proportions in the bulk samples, which is in turn are used to deconvolute cell-type specific effect from the bulk data. A reference panel can be created through expression profiling of sorted cells. However, while good nuclear protein markers exist for sorting nuclei into broad groups of neurons and glia, there is a lack of known, high fidelity, antigens and antibodies for further sorting subclasses of these brain cells. A better alternative is therefore to create the reference panel from s_n_RNA-seq data^15^ that allows a fine grained analysis of brain cell-types.

Even with the advent of s_n_RNA-seq, deconvolution is likely to remain pertinent for association studies with brain tissue. First, the vast majority of existing gene expression data sets involves bulk samples. Deconvolution allows the (re-)use of this “legacy” data to study cell-type specific effects. Second, once the cell-type proportions are estimated, any bulk brain data can be deconvoluted including transcript level expression data, different types of RNA data (e.g., microRNA), or epigenetic data^16^. In contrast, commonly used s_n_RNA-seq protocols are limited to the study of mRNAs with poly-A tail (i.e., protein coding genes and certain long non-coding RNAs). Furthermore, s_n_RNA-seq studies are typically performed on a gene level. If only specific transcripts of the gene are differentially expressed, this will weaken association signals and result in a loss of potentially critical biological information. Third, bulk expression data involves cytoplasm RNA from whole cells that may contain transcripts not present in the nucleus^17,18^. Thus, deconvolution may give a more complete picture of differentially expressed genes. Fourth, deconvolution is potentially useful to validate findings from s_n_RNA-seq studies. Validation with a different technology can eliminate possible false discoveries due to s_n_RNA-seq specific technical artefacts and therefore allows for more rigorous conclusions.

In this study we create a novel reference panel by combining data from the seven largest published s_n_RNA-seq studies in human post-mortem brain samples^19-23^. All brain samples were from the prefrontal cortex, a brain region of key importance for higher level brain processes such as cognition, emotion, and memory. To estimate the cell-type proportions, needed to deconvolute cell-type specific effects from bulk data, we propose an estimator that is suitable for a fine grained analysis of brain cell-types including multiple low abundant cells. Furthermore, to avoid the use of a very large data set and prevent challenges with public access of nuclei level data, our estimator uses a panel comprising mean expression levels rather than nuclei level s_n_RNA-seq data. Finally, we study how to best use this fine-grained panel to optimize power and avoid false discoveries in empirical transcriptome-wide association studies with bulk data.

## METHOD

This section summarizes the methods. Details are given in the supplemental material (e.g., S1.1 refers to section 1.1 in the supplemental material).

### s_n_RNA-seq data sets, quality control and data processing

We downloaded FASTQ files from seven published s_n_RNA-seq in post-mortem brain samples^19-23^. All brain regions involved the prefrontal cortex, predominantly from Brodmann areas BA6, BA8, BA9, BA10, and BA24. To avoid confounding the expression values in the panel by disease processes or disease specific cell states, only the unaffected “controls” from these datasets were used in our study.

All included studies had partitioned nuclei using the Chromium Controller (10X Genomics) and sequenced their libraries using sequencing platforms from Illumina. We used the cellranger^24^ software for aligning the reads to GRCh38 and creating a matrix of unique molecular identified (UMI) counts (i.e., the number of unique molecules for each gene detected in each nucleus). s_n_RNA-seq data primarily yields reads derived from mature spliced RNA (mRNA), which maps to exonic regions but may also capture unspliced pre-mRNA transcripts that can generate intronic reads^25-27^. As nuclei contain a relatively large fraction of pre-mRNA molecules and such molecules are particularly abundant in brain tissue^28^, to obtain a comprehensive picture of gene expression we counted intronic reads as well^29^.

Next, we performed quality control (QC) on samples and nuclei using exactly the same criteria across all studies. Specifically, we eliminated samples with very high levels of debris (Figure S3). In addition, we removed nuclei with very low (indicating low-quality nuclei or empty droplets) or high (indicating “multiplets” that capture expression levels of multiple nuclei) gene and UMI counts (Figures S2 and S3). Finally, nuclei with a high percentage of reads mapping to mitochondrial genes (possible indicating artifacts stemming from sample preparation) were eliminated.

After QC, for each study separately the count data was log-normalized to obtain more normal distributions and reduce effects of possible outliers. Furthermore, genes were given equal weight by scaling the log-normalized count data to have a mean of zero and a standard deviation of one to avoid that highly expressed genes dominate the cluster analyses.

### Clustering

To identify cell-types and cell-states, we performed a cluster analysis in Seurat^30^ (S.1.1). We used analyses that “anchor” the different datasets in a shared cluster space to facilitate integration^31^. The cluster analysis was limited to the 2,000 genes that exhibited the highest nucleus-to-nucleus variation (i.e., highly expressed in some nuclei and lowly expressed in others)^32^. There are potentially a large number of donor-level variables (e.g., sex, age) and covariates (e.g., cDNA yield, post-mortem interval, tissue pH levels, percentage of reads aligned) that may obscure the separation of clusters. To remove this donor-level variation, we regressed out “dummy” variables that indicated the individual samples. Furthermore, we regressed out nuclei related QC indices (e.g., number of genes per nucleus, UMI counts per nucleus cell, and percentage of reads mapping to mitochondrial genes).

### Deconvolution

Deconvolution involves three steps. First, a reference panel^33,34^ is created (S1.2). To select genes for the panel, we used MAST^35^ that performs significance tests to identify the genes that best discriminate between the cell-types. The expression values from the s_n_RNA-seq data were scaled to have a mean of zero and variance of one for each study, and then an average expression value was computed across all studies.

Second, the reference panel in combination with the bulk RNA-seq data is used to estimate cell-type proportions in each bulk sample. To avoid working with a very large data set and prevent challenges with public access of nuclei level data, our estimator uses a panel comprising mean expression levels rather than nuclei level s_n_RNA-seq data. Specifically, we use the standard linear model^36^ but estimated by empirical Bayes^37^ that is more suitable for a fine grained analysis of brain cell-types that may involve multiple low abundant cells. The mean and twice the standard deviation of estimates produced by fitting the same model subject to a non-negativity constraint for the regression coefficients (i.e., the cell-type proportions) was used as the prior distribution.

Third, the estimated cell-type proportions are used to perform cell-type specific association studies with bulk data, This was done by fitting, for each transcript, the deconvolution model described elsewhere^13^ (S1.3). These analyses were performed using the Bioconductor package RaMWAS^38^.

### Demonstration bulk RNA-seq dataset

Bulk RNA-seq data was generated using tissue from BA10 of from 291 control individuals and 304 individuals that were diagnosed with a psychiatric disorder (S1.4). The RNA-seq data was generated using the TruSeq Stranded Total RNA library kit. The sequenced reads were aligned with HISAT2 (v.2.1.0) and transcriptome assembly was performed with StringTie^39^. All analyses (i.e., cell-type proportion estimation and deconvolution analyses) regressed out the covariates: sex and age, indicator variables to account for possible brain banks effects, and assay-related covariates such as total number of reads and the percentage of reads aligned. Furthermore, to account for remaining unmeasured sources of variation, six principal components (as indicted by the scree plot) that were used as covariates after regressing out the measured covariates from the bulk RNA-seq data.

## RESULTS

### Sample description and QC

In total, the seven datasets included s_n_RNA-seq data from 94 unaffected “control” subjects. The sample comprised 37% females. The mean age was 61.6 years (SD=28.6 years) with the 5^th^/95^th^ percentiles of 12.7/90.0 years indicating a very broad range. The post-mortem interval was 19.6 hours (SD=15; 5^th^/95^th^ percentile of 2.5/49.4 hours).

Table S1 lists assay related statistics. In summary, we obtained an average 65,118 reads per nucleus of which 93.6% mapped to the genome and where 78.8% of reads had nucleus-associated barcodes. Using the same criteria for all seven studies, we quality controlled samples and nuclei (S2.2, Figures S1-S3). Two studies had many more nuclei per donor (34,342 and 22,831 nuclei) than the other five studies (mean 5,154 nuclei). To avoid that the clustering was mainly driven by these two studies, we down-sampled their nuclei to 8,562 and 8,567 to obtain an average of 5,547 nuclei (range 1,426-10,039 nuclei) across all seven studies. After QC and down-sampling, 353,146 nuclei from 92 donors remained.

### Clustering and cell-type labeling

Clustering identified 20 groups of nuclei, 17 of which were observed in all seven studies. The three clusters that were not consistently observed were removed from further analyses. Figure 1 visualizes the cell-type clusters. To plot the clusters, which differ on many dimensions, in a two-dimensional space we used Uniform Manifold Approximation and Projection (UMAP).

**Figure 1.**
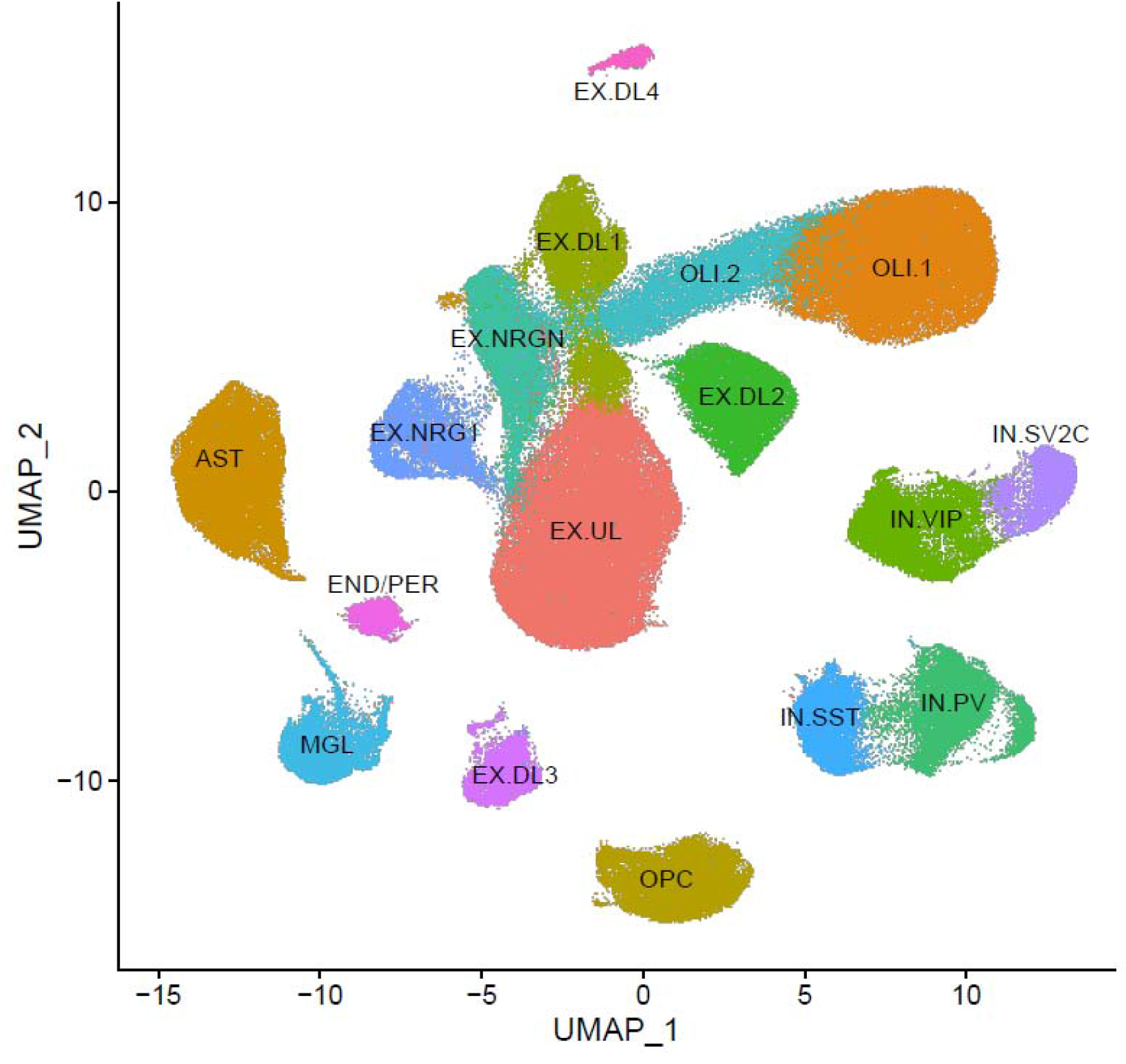
A Uniform Manifold Approximation and Projection (UMAP) plot depicting the nuclei of the 17 clusters. The cluster labels are described in the main text.

Table S2 provides for each cluster a list of standard gene expression markers with high expression levels as well as the most frequently assigned original cell-type label in the five studies that provided labeled nuclei. Of the 17 clusters, 14 could readily be labeled using standard markers. Although it should be noted that only two studies attempted labeling subtypes of broad groups of nuclei (e.g., excitatory neurons), the nuclei of the 14 clusters were consistently labeled by the five studies that provided the original cell-type labels. These 14 clusters included one of the two clusters of oligodendrocytes (OLI.1)^40^, oligodendrocyte precursor cells (OPC)^40^, astrocytes(AST)^41^ and microglia (MGL)^42^. Four clusters of interneurons (IN) were identified that could further be labeled based on the expression of somatostatin (IN.SST), parvalbumin (IN.PV), vasoactive intestinal peptide (IN.VIP), and synaptic vesicle glycoprotein 2C (IN. SV2C)^43^. Finally, seven groups of excitatory neurons were identified. These neurons were further subdivided into one cluster of upper-layer (EX.UL) neurons and four clusters of deep-layer (EX.UL1-EX.UL4) neurons all expressing FOXP2 and subsets of other standard layer-specific markers. Furthermore, we observed neurons expressing neurogranin (EX.NRGN).

Three clusters could not unequivocally be labeled with standard marker and were also inconsistently labeled across the five studies that provided labels for individual nuclei. First, we observed a cluster expressing standard markers for both endothelial cells^44^ and pericytes^37^. In the original studies these nuclei were labeled as endothelial cells^44^, pericytes ^37^, or as a combined cluster of endothelial cells and pericytes. As these nuclei most likely included both endothelial cells and pericytes that have very similar expression profiles relative to the other clusters in Figure 1, were labeled this cluster END/PER.

Second, albeit at relative modest levels compared to OLI.1, the second cluster of oligodendrocytes (OLI.2) expressed standard oligodendrocytes markers MBP, PLP1, and MOBP. In addition, we observed the expression of NRGN, CAMK2A, and CAMK2B that share a motif with MBP potentially allowing it to be packaged together for cytoplasmic transport to dendrites^45^. Three studies labeled these nuclei as oligodendrocytes and the other two studies as neurons. Neurons can use the same packaging mechanism for cytoplasmic transport of the RNAs to dendrites and this potentially explains the confusion about the identity of this second group of oligodendrocytes.

Third, a cluster of EX neurons expressed only few of the markers expressed by the other EX clusters and was inconsistently labeled with respect to cortical layer in the two studies that labeled EX subtypes. This EX cluster expressed NRG1 at very high levels (EX. NRG1). NRG1 is expressed in multiple cell-types and best known as a gene affecting a range of psychiatric and neurological disorders such as Alzheimer, autism and schizophrenia^46,47^. To learn more about the identity of this cluster, we selected the ten most highly expressed genes from the reference panel. Six of the ten genes were previously reported to be associated with a range of psychiatric and neurological disorders. In addition to NRG1^46,47^, this included ZNF804B^48^, CDH12^49^, CLSTN2^50,51^, RIT2^52^, and MCTP1^53^. This pattern is somewhat reminiscent of so-called Von Economo neurons (VENs) that are known to be altered in diseases such as Alzheimer, autism, and schizophrenia^54-56^. VENs are found in humans and great apes (but not other primates), cetaceans, and elephants, and may have evolved for the rapid transmission of crucial social information in very large brains^57^. In humans, VENs are abundant in the anterior cingulate and frontoinsular cortices but are also present in the prefrontal cortex^58^. A recent study involving 879 nuclei from frontoinsula layer 5 identified several VEN markers, but these markers were not highly expressed in our cluster.

### Cell-type proportion estimation

Table S3 shows the MAST^35^ test results identifying 1,652 genes for the reference panel (Table S4). The cell-type estimation procedure was first evaluated using artificial bulk data. We generated artificial bulk data using the cell-type specific expression values from the panel in combination with cell-type proportions that were randomly drawn from a generalized beta distribution assuming the mean, standard deviation minimum, and maximum of the cell-type proportions observed in our demonstration bulk RNA-seq dataset. Table 1 shows that the mean of the estimated cell-type proportions was very close to the cell-type proportions used to simulate the data (correlation is *r*=0.999) and that the estimates were unbiased with a small root mean squared error. Furthermore, only very few of the cell-type proportions were estimated at zero, which indicates that cell-types proportions can be estimated precisely and do not degrade if they are rare.

**Table 1.**
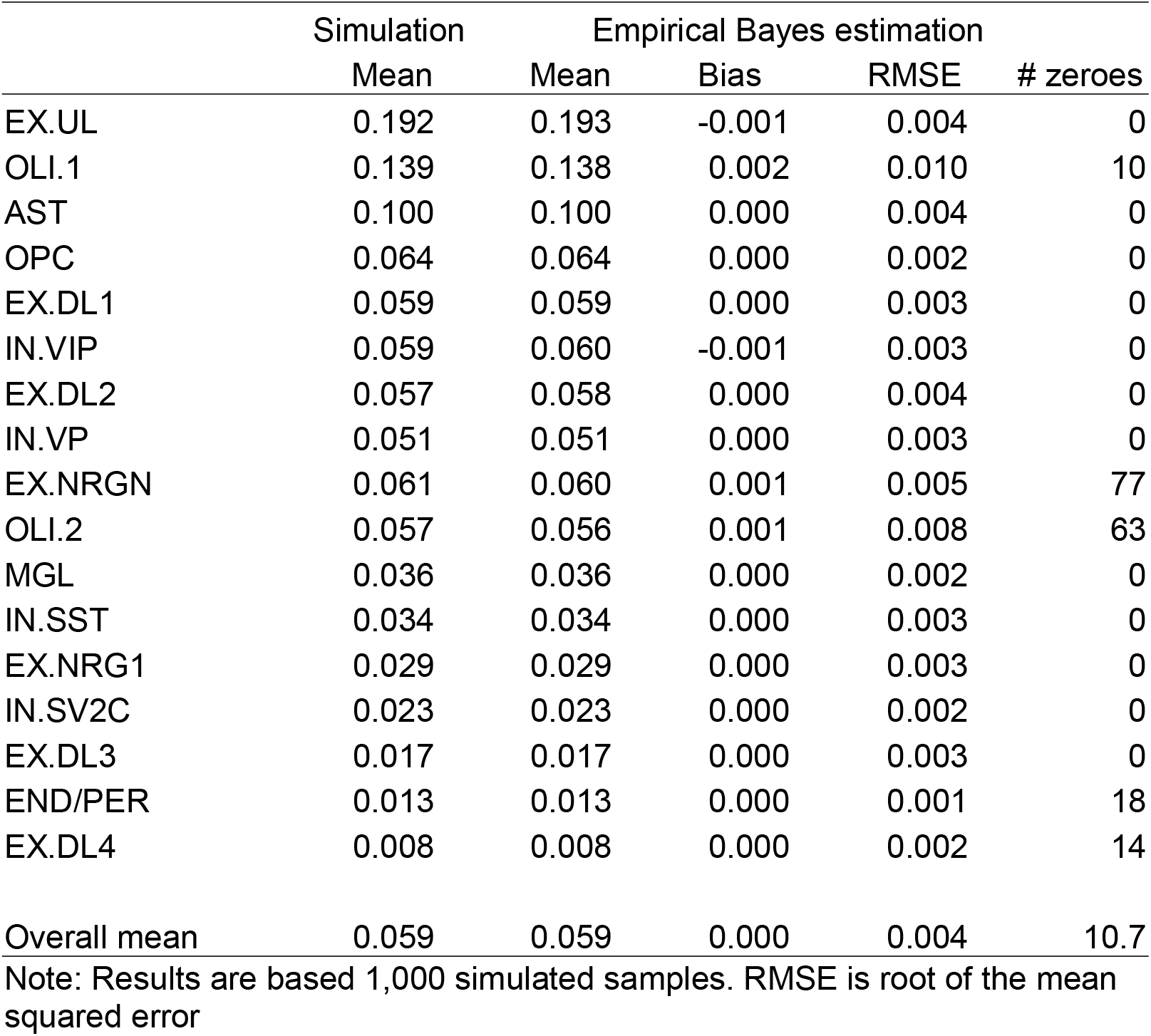
Simulation study to evaluate the empirical Bayes estimation procedure of cell-type proportions

Figure 2 shows that the mean of the cell-type proportion estimates in our demonstration bulk RNA-seq dataset was highly correlated with the mean s_n_RNA-seq counts (*r*=0.994). Only EX.NRGN showed a notable difference. Given that our simulation study yielded unbiased estimates, this most likely reflects true biological variation between the two sample sets. As we observed in the simulation study, the estimates had other favorable properties such as that even for rare cell-types few estimates were estimated to be zero (average 3.2%).

**Figure 2.**
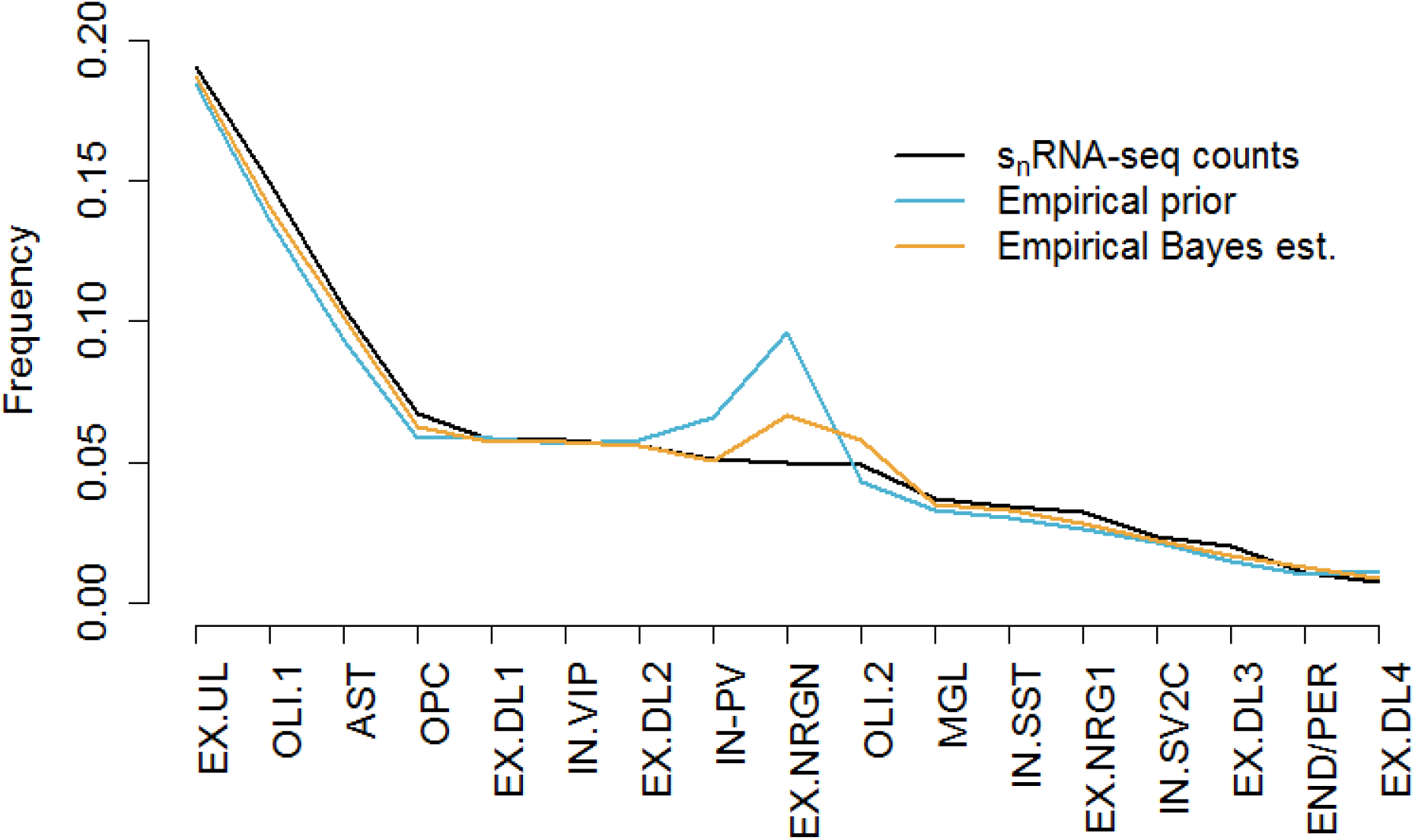
Mean cell-type frequencies observed in the s_n_RNA-seq data and estimated in bulk samples.

The empirical Bayes estimates were slightly better than the estimates used to create its prior distribution. Both estimates were substantially better than the standard ordinary least squares approach^36^ that yielded a lower correlation with the mean s_n_RNA-seq counts (*r*=0.886) and estimated 1.4 times more cell-type proportions to be zero. We further explored whether the estimates could be improved by using the nuclei level s_n_RNA-seq data rather than a panel of mean expression values. For this we used the MuSiC^34^ package that is specifically designed to work with a panel of s_n_RNA-seq data. We had to down-sample the number of nuclei to avoid excessive run times. We could not replicate our previous observation that MuSiC produces superior estimates^59^. In fact, correlations with s_n_RNA-seq counts were very poor (*r*=0.201) possibly indicating convergence problems or challenges combining data from multiple studies.

### Deconvolution

To study whether the distribution of the tests statistics under the null hypothesis followed the assumed theoretical distribution, 1,000 transcriptome-wide association studies (TWASs) where performed after randomly permuting case-control labels. Results showed that the mean/median lambda (ratio of the median of the observed distribution of the test statistic to the expected median) was close to one (Figure 3). This implied the absence of test statistic inflation and that under the null distribution accurate P values are obtained. This was true for even the rarest cell-types suggesting that it is possible to perform TWASs on rare cell-types without an increased risk of false positives. All lambdas obtained from TWASs of observed data were within the 95% confidence intervals of the lambdas observed in the permuted datasets. Although this strictly speaking does not have to be the case, this also suggests the absence of test statistic inflation.

**Figure 3.**
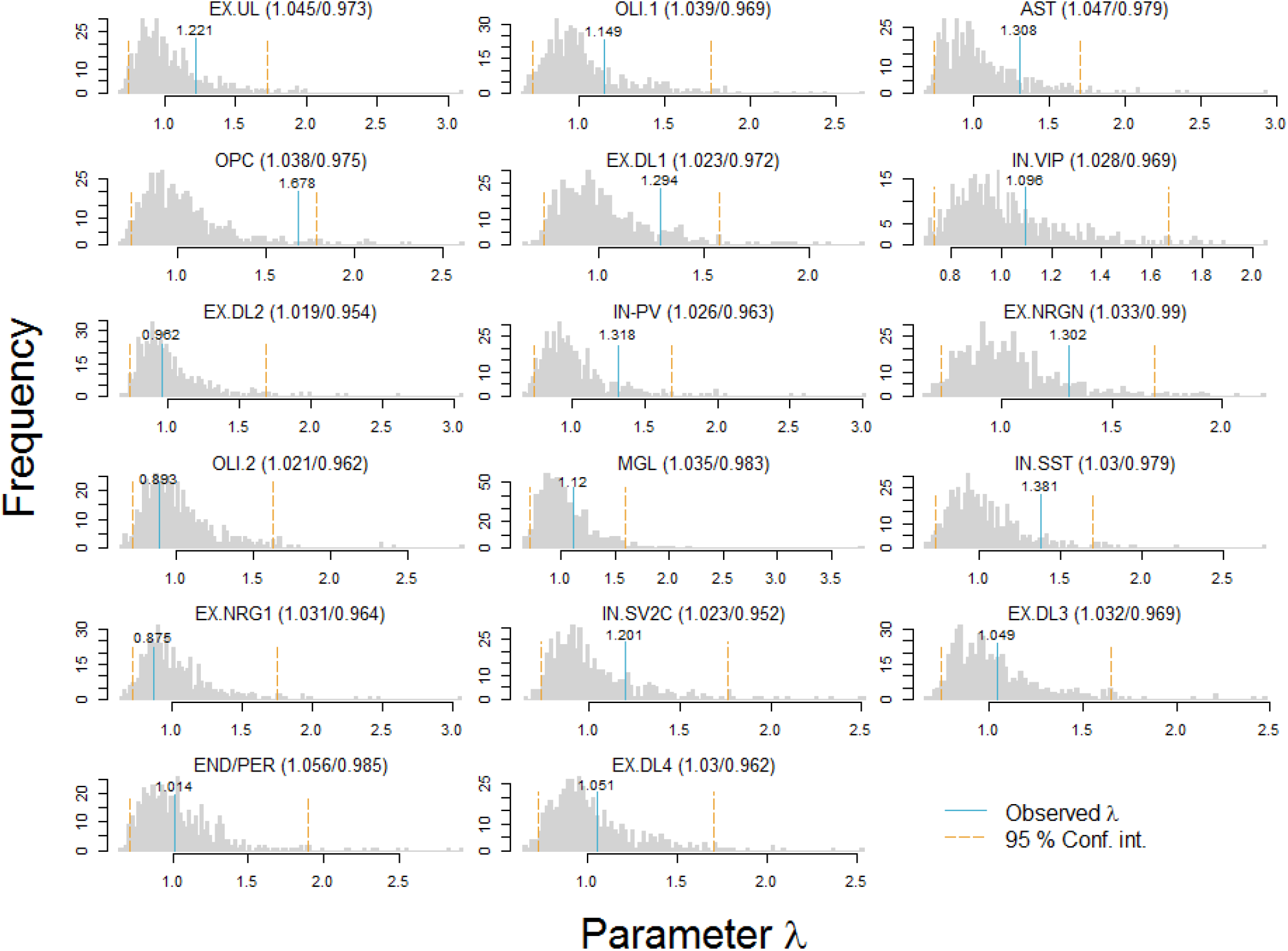
Histograms of observed and simulated lambdas.

In the TWAS of the observed data, the number of significant results was positively correlated with the cell-type proportions. This is most likely the result of lower power due to a “restriction of range” as variation in low abundant cell-types is restricted at zero. We studied whether power for these rare cell-types could be improved by grouping them with other cell-types. For this purpose, we performed a principal components analysis (PCA) followed by varimax rotation on the gene expression values of the panel. In addition, we studied whether omitting rare cell-types altogether improved power for the other cell-types as fewer parameters are estimated.

The PCA suggested that twelve cell-types could be combined in six groups of two cell-types (Table S5). These PCA results corresponded very well with the UMAP plot (Figure 1) showing proximity of these same sets of groups. With one exception (e.g., the combined group EX.UL/EX.NRG1 showed more significant findings than EX.UL and EX.NRG1 together) grouping resulted in fewer significant findings. The likely reason is a dilution of effects when two cell-types are combined that have association signals in different genes (e.g., OLI.1 versus OLI.2 and IN.SST versus IN.PV). Furthermore, we observed that combining/omitting cell-types reduced the number of significant findings of cell-types that were not grouped themselves (e.g., AST, MGL). The likely reason is that grouping cell-types reduces the overall explained variance of the regression model used to perform the cell-type specific TWASs (S1.3) thereby lowering power to detect effects. Overall, these results suggest that best strategy is not to group/omit cell-types.

## DISCUSSION

We created a novel reference panel that allows the detection of differentially expressed genes in human bulk brain data on a fine-grained cell-type specific level. To create the reference panel we analyzed data from the seven largest publicly available s_n_RNA-seq studies. We selected the 17 cell-types for the panel that were robustly detected across all studies. In addition to many known cell-types we detected a group of neurons (EX. NRG1) of potential specific importance to psychiatric and neurological disorders.

To estimate the cell-type proportions, we proposed an empirical Bayes estimator that yielded highly accurate and unbiased cell-type proportion estimates even for the low abundant cell-types. Furthermore, to avoid the use of a very large dataset and prevent challenges with public access of nuclei level data, our estimator has the desirable property that it uses a panel comprising mean expression levels rather than the nuclei level s_n_RNA-seq data. Whereas the panel was created using nuclear mRNA, our bulk mRNA data assayed cytoplasm RNA from whole cells that may contain transcripts not present in the nucleus^17,18^. However, the mean of the cell-type proportion estimates in our bulk RNA-seq dataset was almost identical to the mean cell-type proportions observed in the s_n_RNA-seq data (*r*=0.994, Figure 2). This suggested that any possible differences in expression levels of the genes in the panel in the nucleus versus cytoplasm did not distort the estimates.

Transcriptome-wide association studies performed with permuted bulk RNA-seq data showed that it is possible to perform TWASs for even the rarest cell-types without an increased risk of false positives. Furthermore, the best strategy to optimize power analyze all cell-types in the panel and avoid grouping or omitting (rare) cell-types. For example, even cell-types with frequencies as low as 1% yielded transcriptome-wide significant results in the absence of test statistic inflation.

The proposed approach requires bulk expression data to estimate the cell-type proportions. The estimated cell-type proportions can subsequently be used to deconvolute cell-type effects from the bulk expression data but also any other bulk data generated for the same brain samples (e.g., methylation data, open chromatin data). In this sense the method is generic and not limited to expression data.

In summary, brain disorders are leading causes of disability world-wide. We proposed a novel reference panel and tool set that allows the use of bulk brain data to study brain disorders on a fine-grained cell-type specific level. When studying bulk brain data, the use of this approach may prevent that many disease associations remain undetected. Furthermore, identifying the specific cell-types from which association signals originate is key to formulating refined hypotheses about the etiology of brain disorders, designing proper follow-up experiments and, eventually, developing novel clinical interventions. The reference panel, consisting of profiles from 17 unique brain cell-types, and the accompanying, easy to use, analysis tools are publicly available.

## ACKNOWLEDGEMENTS

This research was supported by grants R01MH109525 and R01MH124981 from the National Institute of Mental Health. The sponsors had no role in the design and conduct of the study; collection, management, analysis, and interpretation of the data; preparation, review, or approval of the manuscript; or decision to submit the manuscript for publication.

## AVAILABILITY OF DATA AND MATERIALS

The panel and R scripts used for empirical Bayes estimation and its evaluation though simulations are available from GitHub: https://github.com/ejvandenoord/Empirical-Bayes-estimation-of-cell-type-proportions. RaMWAS is freely available from Bioconductor (https://bioconductor.org/packages/release/bioc/html/ramwas.html) and a script to deconvolute cell-type effects from bulk data with RaMWAS is also provided on GitHub https://github.com/ejvandenoord/celltype_MWAS

## CONTRIBUTIONS

KA and EO conceived the research question. EO performed the analyses and wrote the first draft of the article. Both authors edited the article and approved the final draft.

