## Supplemental material for "Fine-grained deconvolution of cell-type effects from human bulk brain data using a large single-nucleus RNA sequencing based reference panel"

### S1 METHODS

##### S1.1 Clustering

We used Seurat(1) to identify clusters of nuclei with similar expression profiles. We used integrative analyses(2) that first identifies matching nuclei between the groups called ‘anchors’ followed by clustering all nuclei in the shared space defined by the anchors. To improve results from cluster analyses(3), we limited analyses to the 2,000 genes that exhibited the highest nucleus-to-nucleus variation (i.e., highly expressed in some nuclei and lowly expressed in others). Donor-level variation (e.g., due to demographic variables) and confounders (e.g., cDNA yield, percentage of reads aligned) may obscure the separation of clusters. We therefore regressed out (i) “dummy” variables that indicated the individual samples from the individual donors, and (ii) the QC measures discussed below (S2.2, i.e., log 10 number of genes per nucleus, log 10 UMI counts per nucleus, and % of reads mapping to ribosomal genes). Next, we constructed a K-nearest neighbor graph based on the Euclidean distance of the space defined by the 15 principal components (PCs) that explained most of the variation in the data, and further refined the edge weights between any two nuclei based on the shared overlap in their local neighborhoods. The resolution parameter was set to 0.2 as that gave a robust set of well-delineated clusters. Uniform Manifold Approximation and Projection (UMAP) was used to visualize the cell-type clusters in two-dimensional space. Compared to linear techniques, UMAP may provide a clearer picture of the clusters as nuclei that are close to one another in the original higher-dimensional space are more likely to remain be close to one another in the low two-dimensional space.

##### S1.2 Creating the reference panel

MAST(4) was used to select only the most informative genes for the panel that best discriminate between the identified clusters. Prior to creating the panel, we eliminated nuclei that were outliers with respect for the cell-type they were assigned to. For this purpose, we calculated for each nucleus an outlier score that was the mean of the absolute differences between all the feature means for that nucleus and the corresponding feature means across all nuclei of that cell-type. Next, we calculated the median absolute deviation (MAD) of these outlier scores and identified outliers as nuclei with absolute( outlier score - median( outlier score)) / MAD( outlier score) > 3. We use the MAD rather than standard deviations to define outlier as the MAD is the more robust measure(5).

##### S1.3 Deconvolution of cell-type effects from bulk data

The statistical model for the cell-type specific analyses is:

$$Y^{bulk}= \sum_{c=1}^{n_{c}} m_{c}P_{c}+ \sum_{c=1}^{n_{c}} m_{c}^{D}\left( D \times P_{c} \right)+E$$

Thus, measurements from bulk tissue $Y^{bulk}$ are regressed on $c=1$ to $n_{c}$, cell-type proportions $P_{c}$, and the product of disease status (D) and cell-type proportions $(D \times P_{c})$. The model allows for covariates (not shown) and residual effects $E$. Coefficient $m_{c}$ is the effect of cell-type $c$. The case-control difference $m_{c}^{D}$ for cell-type $c$ is used to test the null hypothesis that cell-type means are equal for cases and controls. With the exception of cell-type proportions the same covariates as described for the bulk analyses were included in the model. Note as $\sum_{c=1}^{n_{c}} P_{c}\cong1$, the model does not require a constant. However, occasionally the model is written with a constant, whereby one of the cell-type proportions is omitted(6), which produces identical results(7-9).

##### S1.4 Demonstration bulk RNA-seq dataset

Bulk RNA-seq data was generated from the prefrontal cortex (Brodmann area 10) of 304 cases with a psychiatric disease and 291 controls. High quality RNA (RNA integrity number mean = 8.69, SD = 0.9590) was extracted from ~30mg of brain tissue using the AllPrep DNA/RNA extraction kit (Qiagen). The RNA-seq data was generated using the TruSeq Stranded Total RNA library kit with Illumina Ribo-Zero Plus with 0.7ug of total RNA from each sample as starting material. With the exception of that each reaction was scaled down, 0.7x the standard reaction volume was used, the vendors protocol was followed. In short, for each sample, ribosomal RNA was depleted from high integrity total RNA, cDNA was synthesized and indexed libraries were created. Up to 42 libraries were pooled in equal molarity and sequenced with paired end reads using a 2x150bp sequence configuration on a NovaSeq 6000 instrument (Illumina).

The sequenced reads were aligned with HISAT2 (v.2.1.0), files were processed by Samtools (10) and transcriptome assembly was performed with StringTie (v.1.3.3) (11) using the human reference genome GRCh37 from ENSEMBL. Following the initial transcriptome assembly with the reference genome, the StringTie merge option was used and the assembly was recreated using only the observed transcripts. In other words, all originally assembled transcriptomes, across all samples, were merged to create a project specific transcriptome including all transcript present in the investigated samples. Next, the abundance levels of all transcripts, for all samples, were re-quantified (stringtie -eB) to ensure that expression measures for specific transcripts are comparable across all samples.

All analyses included the covariates: sex and age, indicator variables to account for possible batch effects, and assay-related covariates such as total number of reads and the percentage of reads aligned. Furthermore, to account for remaining unmeasured sources of variation, 6 principal components that were obtained after regressing out the measured covariates from the bulk RNA-seq data, were included as covariates.

### S2 RESULTS

##### S2.1 Alignment and quality control (QC)

We aligned reads with the “include-introns” option in cellranger that uses a gene transfer format (GTF) file that allows for intronic alignments. Table S1 provides sequencing statistics.

##### Table S1. Sequencing statistics (separate file)

Sample QC: Barcode rank plots plot the total barcode count (y-axis) against the rank of each barcode (x-axis) where the highest ranks have the largest totals. Barcodes for nuclei will have significantly more counts associated with them than the barcodes of background “noise”. A steep drop-off is therefore indicative of good separation between the cell-associated barcodes and noise-associated barcodes (e.g., Figure S1b). Conversely, a lack of steep drop-off may indicate low sample quality and many noise-associated barcodes (e.g., Figure S1b). To quantify, we used the fraction of reads associated with nuclei. Ten samples were omitted as less than 50% of the reads were associated with nuclei. This left 704,260 nuclei. For 8 of the 10 samples multiple libraries were available. For 2 of the 10 samples there was only one library so that the total number of controls decreased from 94 to 92 after this QC step.

##### Figure S1 Examples of barcode rank plots

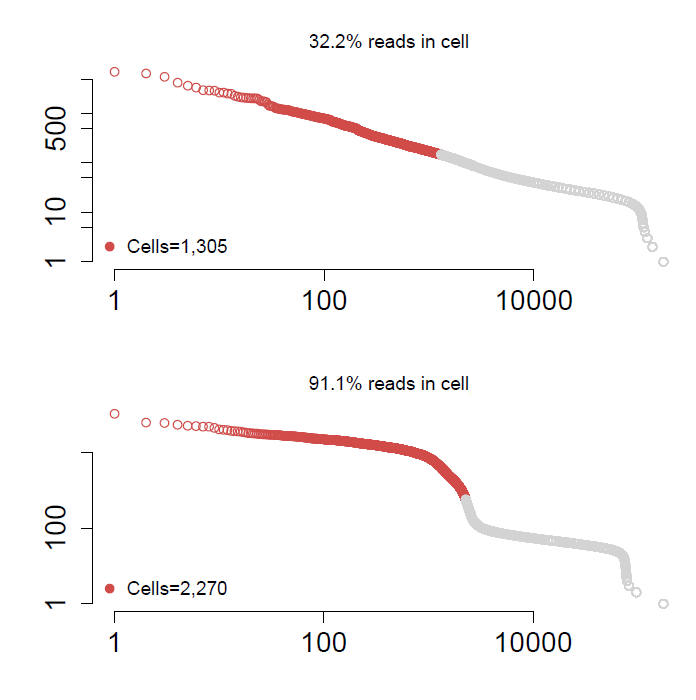

Nuclei QC: Two studies had many more nuclei than the others (309,079 and 182,645 versus the mean of 47,889 for the other 5 studies). To avoid that the clustering will be mainly be driven by these two studies we downsampled their nuclei by randomly selecting sequencing lanes (leaving 77,058 and 58,692 of the 309,079 and 182,645 nuclei). A total of 373,033 nuclei remained. After this QC step, the mean number of nuclei across the 7 studies was 55,005, SD=19,495, range 21,906-77,058).

Low-quality nuclei or empty droplets are likely to have few genes expressed and a small number of UMI counts, whereas nuclei multiplets are likely to have a high gene and UMI count as they capture expression levels of multiple nuclei. We removed 14,021 nuclei with fewer than 400 genes and more than 9,000 genes. These thresholds were chosen as they defined the extreme values of the distribution of number of genes per nucleus across the 7 studies(Figure S2, we used the log base 10 as this transformation was also used prior to performing cluster and association analyses). Next, we eliminated 3,447 nuclei with UMI counts < 500 and with UMI counts > 30,000 (Figure S3). Finally, we removed 2,419 nuclei with more than 5% of reads mapping to ribosomal genes as that may be an artifact stemming from sample preparation. This left 373,033-(14,021+3,447+2,419) = 353,146 nuclei for the cluster analyses.

##### Figure S2. Distribution and QC threshold for number of genes per nucleus

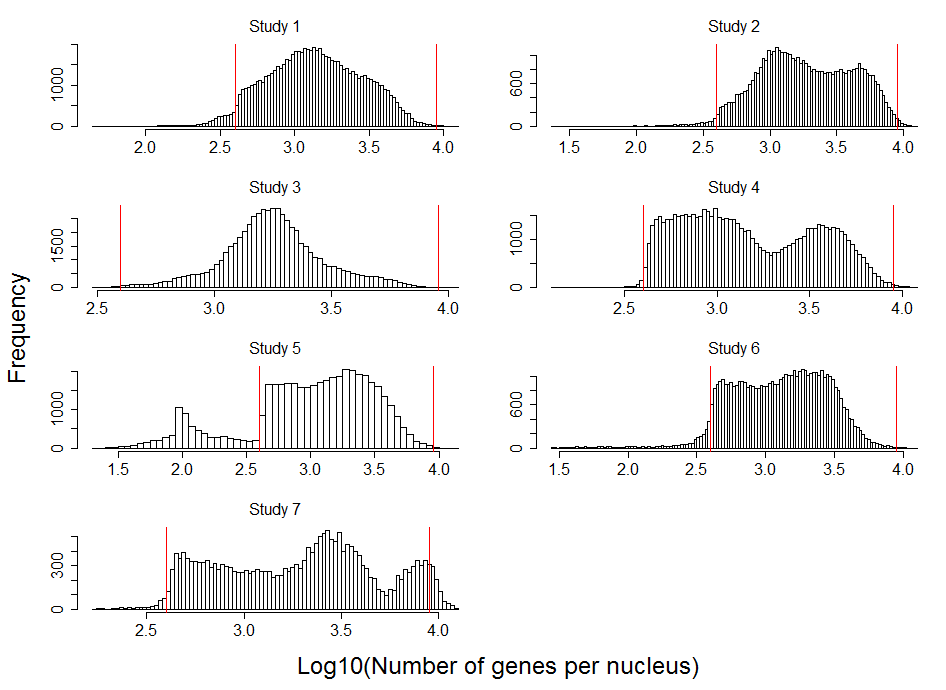

##### Figure S3. Distribution and QC threshold for number of UMI counts per nucleus

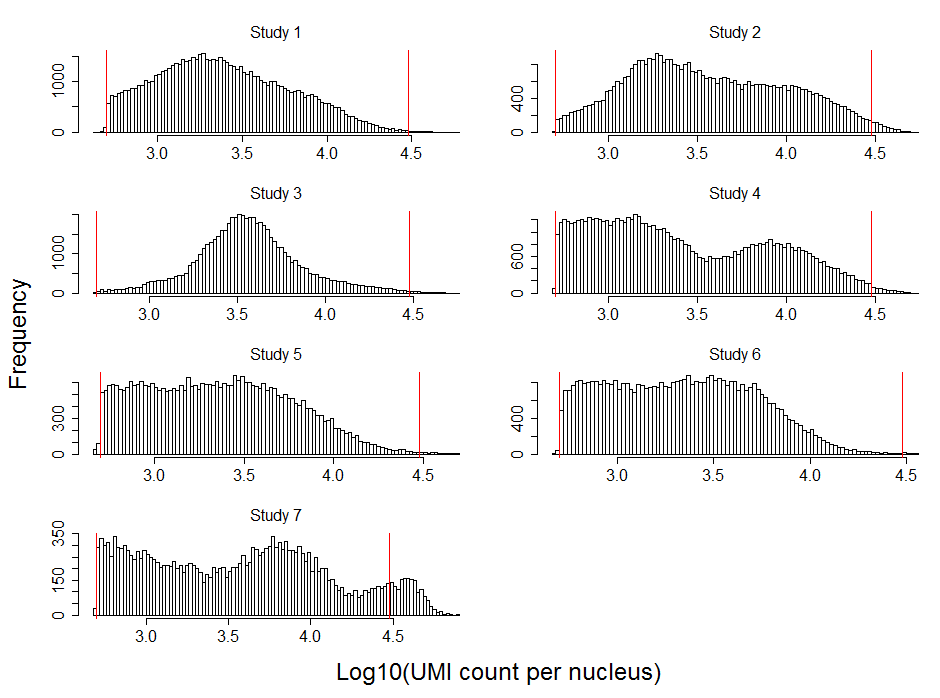

Gene QC: Our data comprised 36,601 genes. For the cluster analyses the 2,000 most highly variable genes were selected (see section S1.4).

Data transformations: UMI count data were log-normalized to obtain more normal distributions and reduce effects of possible outliers. Next, to give equal weight and avoid that highly-expressed genes dominate the cluster analyses, the data was scaled to have a mean expression across nuclei of zero and a variance of one.

##### S2.2 Cell-type identification and labeling

##### Table S2. Labeling cluster (separate file)

##### Table S3. MAST identified cell-type panel markers (separate file)

##### Table S4. Cerebral cortex reference panel (separate file)

##### Table S5. Grouping cell-type by principal components analysis

|  | PC1 | PC2 | PC3 | PC4 | PC5 | PC6 | PC7 | PC8 | PC9 | PC10 |
| --- | --- | --- | --- | --- | --- | --- | --- | --- | --- | --- |
| EX.UL | -0.54 | 0.041 | 0.045 | 0.118 | -0.07 | -0.13 | 0.083 | 0.023 | 0.054 | -0.01 |
| OLI.1 | 0.106 | 0.056 | 0.048 | -0.61 | -0.03 | -0.01 | -0.02 | 0.013 | 0.053 | 0.074 |
| AST | 0.463 | 0.206 | 0.235 | 0.452 | -0.14 | -0.01 | 0.171 | 0.119 | 0.215 | 0.298 |
| OPC | 0.057 | 0.031 | -0.79 | 0.051 | -0.24 | -0 | 0.019 | 0.002 | 0.029 | 0.046 |
| EX.DL1 | 0.057 | 0.066 | 0.032 | 0.056 | -0.01 | -0.11 | 0.105 | -0.01 | 0.01 | -0.74 |
| IN.VIP | -0.01 | 0.103 | 0.029 | 0.04 | -0 | 0.057 | 0.099 | -0.05 | -0.75 | 0.068 |
| EX.DL2 | -0.03 | -0.01 | 0.025 | 0.067 | -0.02 | 0.105 | -0.06 | 0.028 | 0.069 | -0.51 |
| IN.VP | 0.103 | -0.7 | 0.048 | 0.055 | -0.01 | -0.06 | 0.015 | 0.028 | 0.022 | -0.04 |
| EX.NRGN | 0.049 | 0.043 | 0.029 | 0.058 | -0.02 | -0.01 | -0.96 | 0.007 | 0.014 | 0.022 |
| OLI.2 | 0.076 | 0.039 | 0.043 | -0.62 | -0.03 | -0.01 | 0.077 | 0.028 | 0.052 | 0.069 |
| MGL | 0.056 | 0.022 | 0.086 | 0.033 | 0.861 | -0 | 0.012 | 0.003 | 0.018 | 0.02 |
| IN.SST | -0.01 | -0.66 | -0.01 | 0.023 | -0.01 | 0.056 | 0.016 | -0.02 | -0 | 0.082 |
| EX.NRG1 | -0.52 | 0.062 | 0.037 | 0.06 | -0.01 | 0.081 | -0.02 | 0.024 | 0.016 | 0.027 |
| IN.SV2C | 0.061 | -0.06 | 0.012 | 0.049 | -0.02 | -0.06 | -0.07 | 0.059 | -0.6 | 0.021 |
| EX.DL3 | -0.34 | -0.02 | 0.037 | 0.054 | 0.033 | 0.595 | 0.023 | -0.01 | 0.097 | 0.197 |
| END/PER | 0.047 | 0.012 | 0.018 | 0.038 | -0.01 | -0 | 0.006 | -0.99 | 0.03 | 0.027 |
| EX.DL4 | 0.26 | 0.013 | -0.02 | -0.03 | -0.03 | 0.766 | -0 | 0.01 | -0.06 | -0.2 |

Using an absolute loading of 0.5 as the cut-off, Table S5 suggest relatively high similarity in the expression of the (two) groups of OLIs, EX.DL3 and EX.DL4 excitatory neurons, and EX.DL1 and EX.DL2 excitatory neurons. Furthermore, IN.VIP and IN.SV2C interneurons, IN.SST and IN.PV interneurons, EX.UL and EX.NRG1 excitatory neurons.
